# Benchmarking ultra-high molecular weight DNA preservation methods for long-read and long-range sequencing

**DOI:** 10.1101/2021.07.13.451380

**Authors:** Hollis A. Dahn, Jacquelyn Mountcastle, Jennifer Balacco, Sylke Winkler, Iliana Bista, Anthony D. Schmitt, Olga Vinnere Pettersson, Giulio Formenti, Karen Oliver, Michelle Smith, Wenhua Tan, Anne Kraus, Stephen Mac, Lisa M. Komoroske, Tanya Lama, Andrew J. Crawford, Robert W. Murphy, Samara Brown, Alan F. Scott, Phillip A. Morin, Erich D. Jarvis, Olivier Fedrigo

**Affiliations:** University of Toronto, Toronto, Ontario, Canada; The Rockefeller University, New York, New York, United States; Max Planck Institute of Molecular Cell Biology and Genetics, Dresden, Saxony, Germany; Tree of Life program, Wellcome Sanger Institute, Hinxton, Cambridgeshire, United Kingdom; University of Cambridge, Cambridge, Cambridgeshire, United Kingdom; Arima Genomics, Inc., San Diego, California, United States; National Genomics Infrastructure, SciLifeLab, Uppsala University, Uppsala, Sweden; University of Massachusetts Amherst, Amherst, Massachusetts, United States; Department of Biological Sciences, Universidad de los Andes, Bogotá, 111711, Colombia; Johns Hopkins University, Baltimore, Maryland, United States; Southwest Fisheries Science Center, National Marine Fisheries Service, NOAA, La Jolla, California, United States; Howard Hughes Medical Institute, Chevy Chase, Maryland, United States

**Keywords:** long-read sequencing, genome assembly, tissue preservation, HMW DNA extraction

## Abstract

Studies in vertebrate genomics require sampling from a broad range of tissue types, taxa, and localities. Recent advancements in long-read and long-range genome sequencing have made it possible to produce high-quality chromosome-level genome assemblies for almost any organism. However, adequate tissue preservation for the requisite ultra-high molecular weight DNA (uHMW DNA) remains a major challenge. Here we present a comparative study of preservation methods for field and laboratory tissue sampling, across vertebrate classes and different tissue types. We find that no single method is best for all cases. Instead, the optimal storage and extraction methods vary by taxa, by tissue, and by down-stream application. Therefore, we provide sample preservation guidelines that ensure sufficient DNA integrity and amount required for use with long-read and long-range sequencing technologies across vertebrates. Our best practices generated the uHMW DNA needed for the high-quality reference genomes for Phase 1 of the Vertebrate Genomes Project (VGP), whose ultimate mission is to generate chromosome-level reference genome assemblies of all ∼70,000 extant vertebrate species.

## Introduction

The past two decades have seen genome sequencing become increasingly easy and affordable, driven by advancements in sequencing and computing technologies. Growing accessibility spurred the formation of large-scale consortia, such as the Genome 10K project (G10K), with the goal of generating genome assemblies for many species to enable new scientific discoveries and aid in conservation efforts [1]. However, initial efforts used short read sequencing (< 200 bp), such as Illumina technology, which were later found to often result in genome assemblies that were highly fragmented, incomplete, and plagued with structural inaccuracies [1–3]. Subsequently, G10K initiated the Vertebrate Genomes Project (VGP), with the mission of producing high-quality, near-complete, and error-free genome assemblies of all ∼70,000 extant vertebrate species [4]. By comparing sequencing data types and assembly algorithms, the VGP consortium determined that it was not possible to obtain high-quality reference assemblies at the chromosomal level without the use of long-reads (e.g. > 10 kb), such as Pacific Biosciences, long-range molecules (e.g. > 50 kb), such as 10X Genomics linked reads, or optical mapping (> 150kb) such as with Bionano Genomics, and Hi-C proximity ligation (> 1 Mb), such as with Arima Genomics, all of which can span repeats thousands of base pairs in size [4]. To take full advantage of these new sequencing and assembly methods, molecules of DNA need to be as long as possible.

While long-read and long-range (LR) data simplify and accelerate the assembly, they come with a major challenge: they require large amounts of very high-quality DNA. For short-read technologies, many nucleic acid isolation methods developed over the years, including the standard phenol-chloroform method [5] had been sufficient. LR technologies require relatively pure DNA in the 10 kb to 300 kb range. Additionally, the Hi-C method requires physical cross-linking of contacting DNA regions within the same chromosomes, thus requiring cell nuclei to be intact before processing and isolation of cross-linked DNA [4]. With Hi-C, 3D interactions within chromosomes serve to assemble contigs or short scaffolds into chromosomal-scale scaffolds. For LR technologies, only a few extraction methods are currently able to produce high molecular weight (HMW) DNA ranging from 45 to 150 kb or ultra-high molecular weight (uHMW) DNA which is over 150 kb long. These include bead-based (MagAttract HMW DNA Kit, Qiagen), high□salt [6], and agarose plug methods (Bionano Prep Soft/Fibrous Tissue Protocol, Bionano Genomics) [7]. More recently, a less laborious thermoplastic magnetic disks (Nanobinds) method was developed by Circulomics [8]. Regardless of their capabilities, the performance of HMW and uHMW DNA extraction methods primarily depend on the type of sample and how it was collected, handled, and preserved.

The long-held “gold standard” in tissue preservation for high-quality DNA isolation has been flash-freezing tissues in liquid nitrogen directly after collection, followed by ultra-cold –80°C long-term storage [9–14]. While liquid nitrogen is readily available in most laboratory setups, its limited availability in many fieldwork conditions can be an insurmountable hurdle. Indeed, a large portion of global biodiversity is located far from labs, and sampling such species will require long expeditions under rustic field conditions. Thus, transporting sufficient amounts of liquid nitrogen from the point of collection to the laboratory is often infeasible and the applicability of flash-freezing outside the lab environment is greatly limited [10,13,15]. Additional considerations specific to the studied species exacerbate the challenge of sample collection and preservation. DNA degradation is promoted by enzymes whose concentrations are likely to be tissue-specific and possibly species-specific. Small organisms provide little tissue, and preferred tissue types may be unavailable. Permitting restrictions also vary widely among species and among countries. Yet, methods for field sampling in non-model species for the purposes of LR sequencing remain anecdotal or unsubstantiated, as failed attempts are not published and very few preservation experiments have measured fragment sizes relevant to LR technologies [16,17]. Thus, methods that bridge the gaps between uHMW DNA, the lab, and field conditions still require benchmarking.

Here, we perform a series of benchmarking experiments to assess sample preservation methods under laboratory and simulated field conditions and compare the quality of uHMW DNA obtained. Specifically, we extract uHMW DNA from multiple tissue types of representative vertebrate species, which were collected under various preservation and temperature conditions. For each experimental sample, we evaluate the fragment length, yield, and purity of the uHMW DNA extracted. Based on our findings, we propose a new set of guidelines for tissue preservation, ranging from best to minimally adequate practices for acquiring uHMW DNA from both laboratory and field collected samples, necessary for producing high-quality reference genome assemblies.

## Results

In this study, we used the agarose plug method optimized by Bionano Genomics [7] across all species and preservation methods albeit with small protocol variations for fibrous tissues, soft tissues, and blood. We tested six preservation methods (**Fig. 1**): 1) flash frozen in liquid nitrogen, which served as the ‘gold standard’ and our point of reference; 2) 95% ethanol (EtOH), a long preferred method of field preservation of tissues [10,15,18]; 3) 20–25% dimethyl sulfoxide (DMSO) buffer (see Methods), which has been shown to be very effective at permeating tissues and preserving HMW DNA after long-term storage at ambient temperature [19,20]; 4) RNAlater Stabilization Solution (RNAlater; Invitrogen, Waltham, MA, USA), a commonly used preservative that also facilitates transcriptomics; 5) DNAgard tissue and cells (DNAgard; Biomatrica, San Diego, CA, USA), a commercial preservative designed for stabilizing DNA in tissues at room temperature; and 6) Allprotect Tissue Reagent (Allprotect; Qiagen, Hilden, Germany), another commercial preservative targeting stable room-temperature tissue preservation. We exposed preserved samples to different temperatures (4°C, room temperature, and 37°C) for various durations of time (6 hr to 5 months). We did so with up to 6 tissue types (muscle, blood, ovary, spleen, isolated red blood cells (RBCs), and whole-body) from 6 species representing five vertebrate lineages (a mammal, a bird, two turtles, an amphibian, and a bony fish; **Fig. 1**), for a total of 140 samples (**Table S1**). We assessed the fragment length distribution and DNA yield for each DNA sample. Statistical analyses were performed using linear models that included type of preservative, temperature/time treatment, vertebrate group, and tissue type as variables.

**Figure 1.**
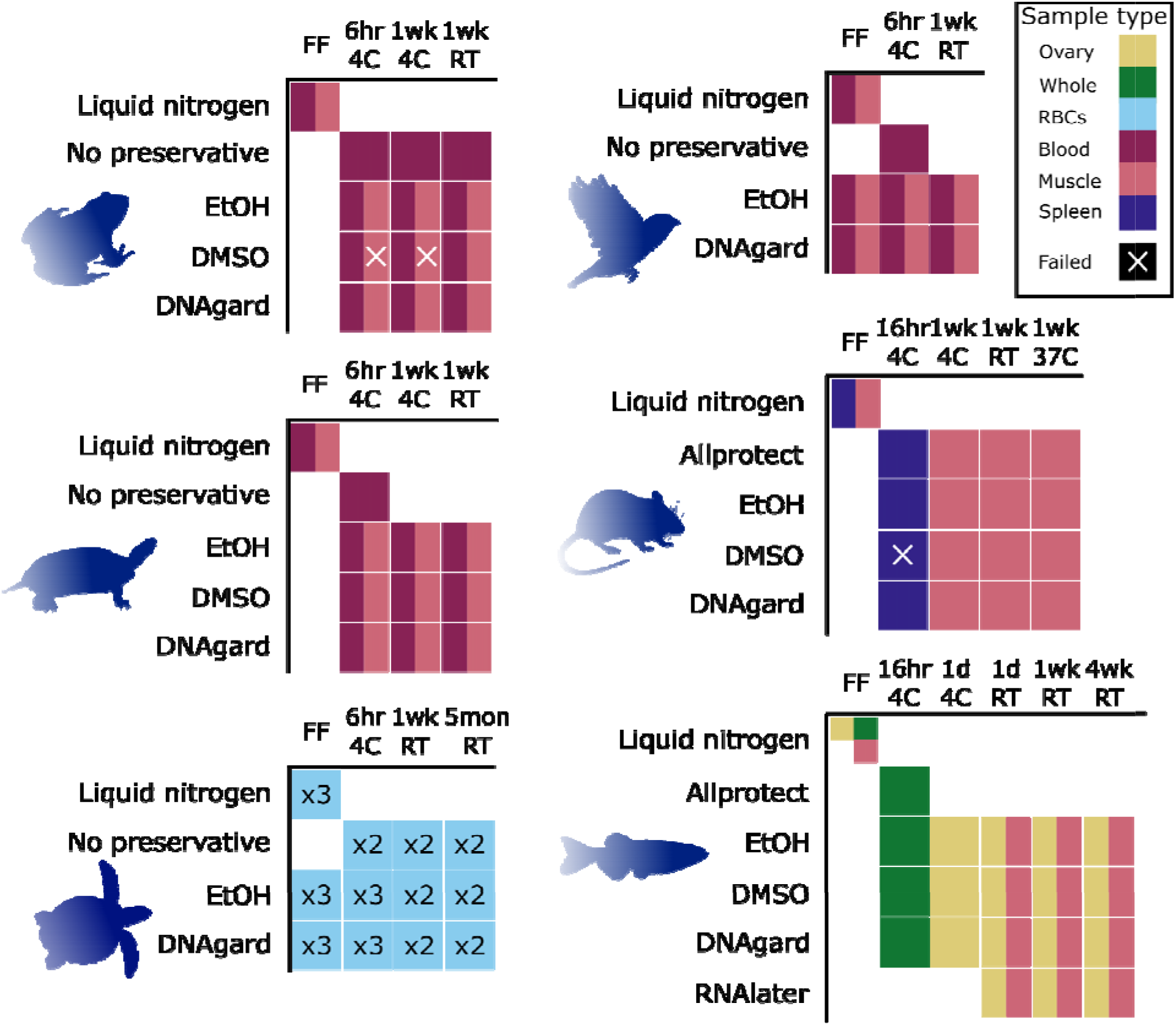
Experimental design for benchmarking tissue preservation. Graphical visualization of samples and treatments used in this study. Rows denote preservative treatments and columns temperature treatments. Colors indicate different types of tissue samples (see legend at top right). All samples were transferred to –80°C after the specified temperature treatment, e.g. ‘6 hr 4C’ means stored at 4°C for 6 hours before transfer to –80°C. Abbreviations are as follows: RBCs, isolated red blood cells; EtOH, 95% ethanol; DMSO, a mix of 20–25% dimethyl sulfoxide, 25% 0.5 M EDTA, and 50–55% H2O; DNAgard, DNAgard tissue and cells cat. no. #62001-046, Biomatrica; Allprotect, Allprotect Tissue Reagent cat. no. 76405, Qiagen); RNAlater, RNAlater Stabilization Solution cat. no. AM7021, Invitrogen; FF, flash-frozen in liquid nitrogen immediately upon dissection; 6hr, six hours; 1d, one day; 1wk, one week; 5mon, five months; RT, room temperature (20–25°C). Samples were collected from these species: house mouse (*Mus musculus*), zebra finch (*Taeniopygia guttata*), Kemp’s Ridley sea turtle (*Lepidochelys kempii*), painted turtle (*Chrysemys picta*), American bullfrog (*Rana catesbeiana*), and zebrafish (*Danio rerio*).

### Fragment length distribution analysis

For extractions that yielded a detectable amount of DNA, we measured their fragment length distributions using at least one of two available techniques: Pulsed-field Gel Electrophoresis (PFGE) and the Agilent Femto Pulse system (FEMTO). PFGE was more informative for analyzing uHMW DNA molecules above 200 kb, due to greater dynamic range in molecular weight separation (**Fig. S1a**), whereas FEMTO was more useful for separating molecules within the 50–165 kb range (**Fig. S1b**). Overall, the agarose plug method yielded high-quality DNA concentrated in the 300–400 kb range (**Fig. 2**).

**Figure 2.**
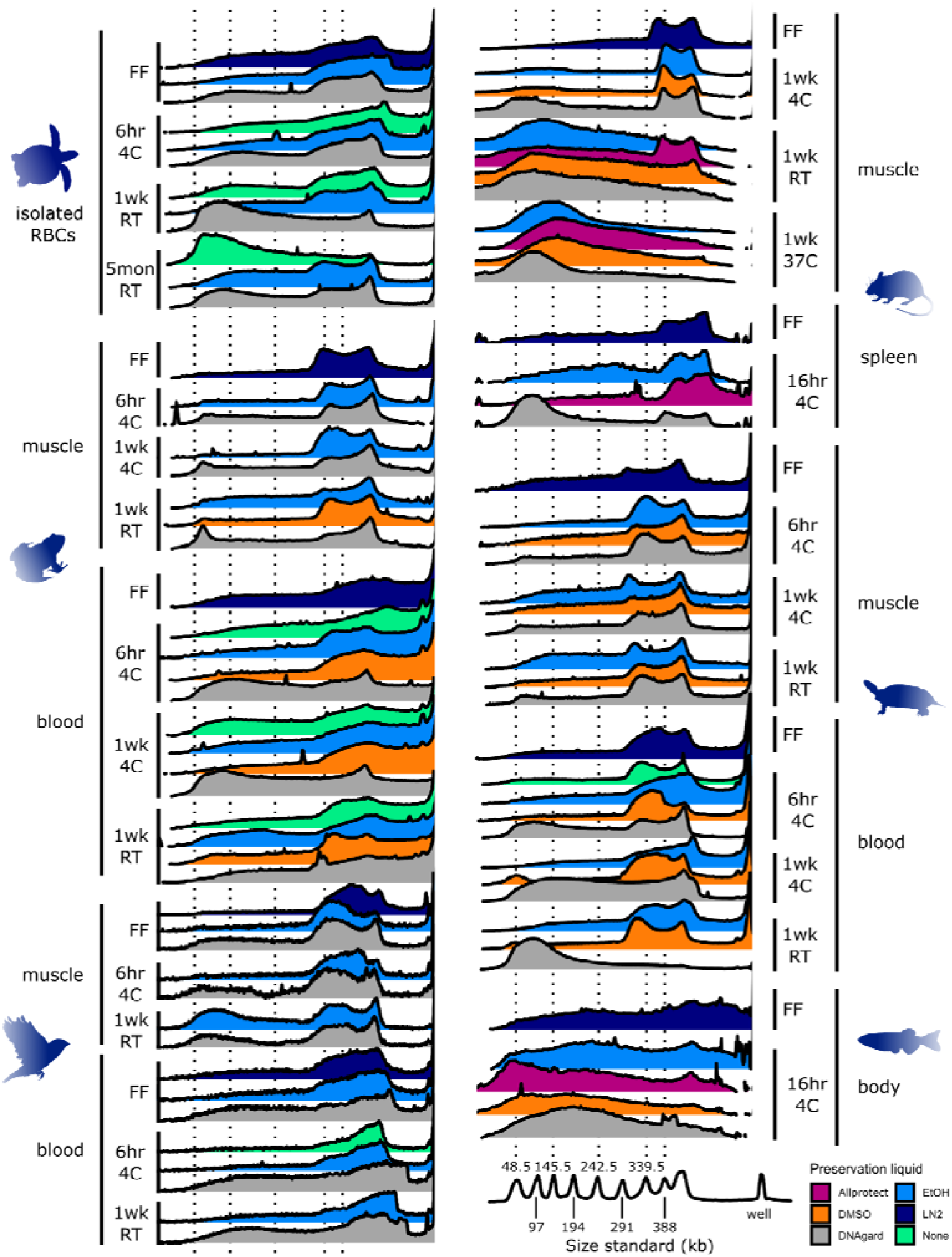
Pulsed-field gel electrophoresis (PFGE) measurements of uHMW DNA comparing different sample temperature and storage times. PFGE traces are visualized as overlapping ridgeline plots. Each ridgeline plot corresponds to a gel lane and a single DNA extract with brightness converted to a plot profile. The x-axis denotes molecule length scaled via piecewise linear scaling to match across gels of different lengths with a common size standard (Lambda PFG Ladder, New England Biolabs). The x-axis is the same in both columns. The y-axis of each plot is a proportional signal in that particular gel lane from just below the well to just beyond the 48.5 kb ladder peak such that the relatively intense brightness of the well itself is excluded. Colors represent different sample preservation methods, as indicated in the legend at bottom right. All samples were transferred to –80°C after the specified temperature treatment, e.g. ‘6hr 4C’ means stored at 4°C for 6 hours before transfer to –80°C. Abbreviations are as follows: RBCs, isolated red blood cells; EtOH, 95% ethanol; DMSO, a mix of 20–25% dimethyl sulfoxide, 25% 0.5 M EDTA, and 50–55% H2O; DNAgard, DNAgard tissue and cells cat. no. #62001-046, Biomatrica; Allprotect, Allprotect Tissue Reagent cat. no. 76405, Qiagen); FF, flash-frozen in liquid nitrogen immediately upon dissection; 6hr, six hours; 1d, one day; 1wk, one week; 5mon, five months; RT, room temperature (20–25°C). Three additional samples were tested, but produced insufficient DNA for fragment length analysis: frog muscle in DMSO for one week at 4°C and 6 hr at 4°C and mouse spleen in DMSO for 16 hr at 4°C. For measurements based on the FEMTO pulse instrument and additional tissue types, see **Figs. S2, S3**.

#### Temperature

From the linear modeling of both PFGE (**Fig. 2, Table S2**) and FEMTO results (**Figs. S2, S3**), we found that temperature treatment was the predictor with the strongest evidence of an effect on the proportion of DNA fragments above 145 kb for PFGE (DF = 6, LR Chisq = 36.62, p = 2.09e-06; **Fig. 3a**) and above 45 kb for FEMTO (DF = 8, LR Chisq = 44.80, p = 4.01e-07; **Fig. S4a**). Samples held at higher temperatures yielded a lower proportion of uHMW DNA, with flash-freezing performing best (**Fig. 3a**). However, samples refrigerated at 4°C for 6 hr following collection were statistically indistinguishable from flash-frozen samples (PFGE: z = 0.56, p = 1.00; FEMTO: z = 2.03, p = 0.48). Samples refrigerated at 4°C for longer periods of up to one week showed some signs of degradation, albeit not consistently across tissue types and species (**Figs. 2, S2, and S3**).

**Figure 3.**
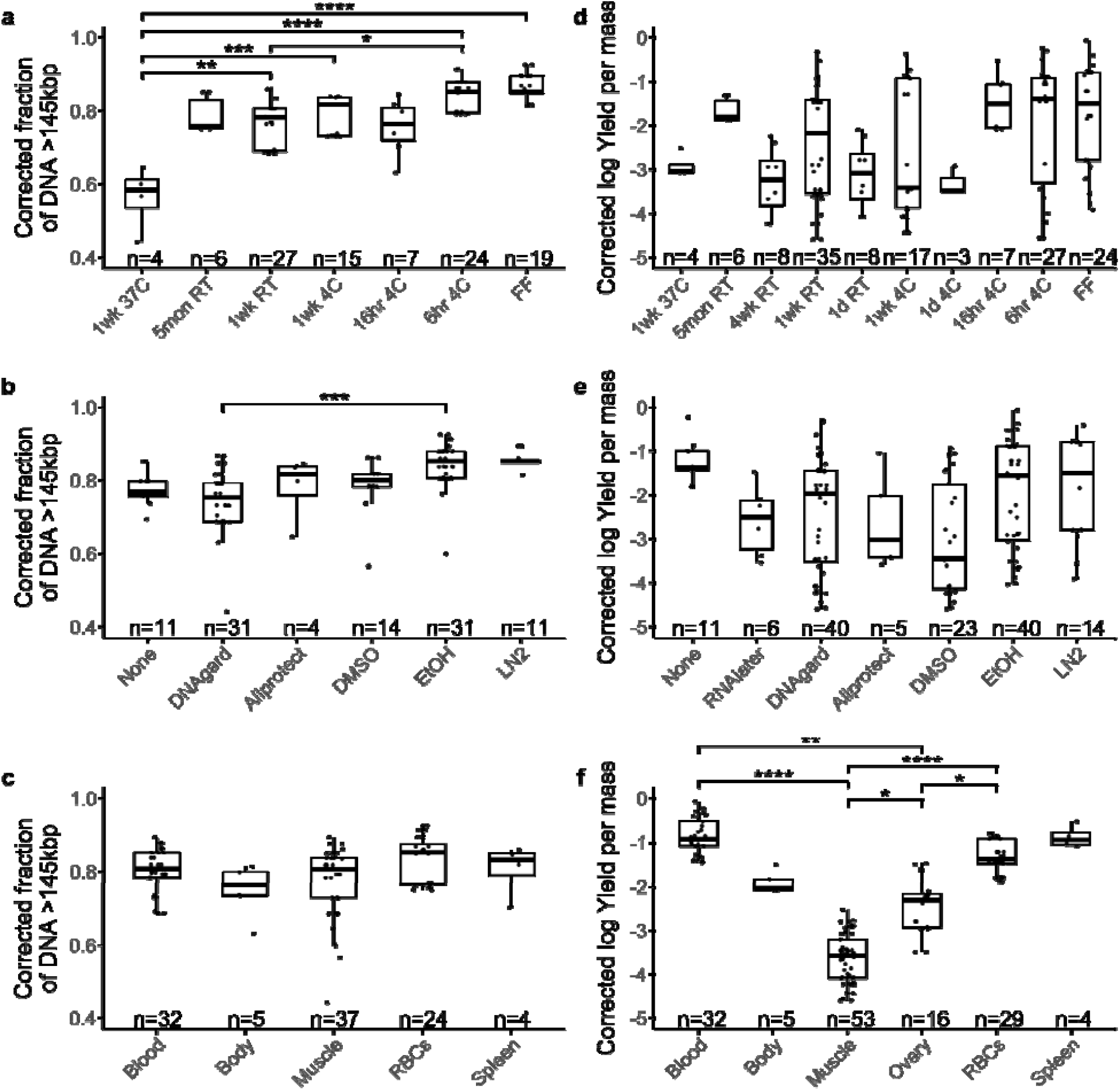
Testing the effect on two measures of uHMW DNA quality. Distributions of sample groups are overlaid with results of linear modeling of fragment length (n = 102, **a-c**) and DNA yield (n = 139, **d-f**). Shown are univariate scatterplots overlain with box plots indicating the median, quartiles, and full range of individual observations. Fragment length was quantified here as the proportion of pulsed-field gel electrophoresis (PFGE) signal above 145 kb, and was modeled in a generalized linear model with temperature (**a**), preservative (**b**), and tissue type (**c**) as predictors. DNA yield per input mass was log-transformed and modeled with temperature (**d**), preservative (**e**), tissue type (**f**), and vertebrate group as predictors. Significant relationships from post-hoc comparisons are shown as connecting bars with significance levels: **** p < 0.0001, *** p < 0.001, ** p < 0.01, * p < 0.05. Sample sizes for each factor are given along the x-axis.

#### Preservation method

The predictor with the second strongest evidence of an effect on the proportion of DNA fragments above 145 kb or 45 kb was preservative treatment (PFGE: DF = 5, LR Chisq = 24.43, p = 0.0002, **Fig. 3b;** or FEMTO: DF = 6, LR Chisq = 25.01, p = 0.0003, **Fig. S4b**, respectively). In PFGE measurements, significant differences were found between DNAgard and EtOH preservation (z = 4.24, p = 0.001, **Fig. 3b**), with DNAgard generally performing poorer. Flash-freezing and EtOH performed better than the other preservation methods in PFGE, and albeit not statistically significant, they had the lowest standard deviation (**Fig. 3b**). Based on PFGE, EtOH was slightly better than DMSO (**Fig. 3b**). Based on FEMTO, DMSO was slightly better than EtOH (**Fig. S4b**). Neither relationship showed significant differences in preservation. In FEMTO measurements, flash-frozen and DMSO-preserved samples showed significantly better preservation efficiency than RNAlater samples (vs. DMSO: z = 3.42, p = 0.009; vs. flash-frozen: z = 3.50, p = 0.007), tested on fish samples. Allprotect outperformed EtOH in room temperature mouse samples but underperformed in the refrigerated fish body set (**Figs. 2 and S3**).

#### Tissue type

Tissue type did not have a significant effect on fragment length overall (**Figs. 3c and S4c, Table S2**). However, muscle showed more variability than blood samples in uHMW DNA yield (> 145 kb). The RBCs samples showed the smallest proportion of degradation, while some muscle samples showed the highest degradation (**Fig. 3c**). In terms of variation between species, the mouse and fish samples showed a higher degree of degradation with respect to temperature treatment than the other species (**Figs. 2, S2, and S3**). It is unclear if this can be explained by a species-specific temperature sensitivity, or if it is caused by technical variation.

#### Interactions among variables

In terms of qualitatively assessing combinations of variables, storage in EtOH appeared to perform best at preserving uHMW DNA for all 4°C refrigerated samples (**Fig. 2**). Notably, nucleated blood samples refrigerated with no added preservatives were stable for up to one week with no substantial signs of degradation (**Fig. 2**). An increased proportion of smaller DNA fragments was evident in refrigerated samples preserved using DNAgard, with the exception of turtle RBCs and muscle samples for which DNAgard results were equivalent to other preservation methods (**Fig. 2**). Fish body samples stored for 16 hr at 4°C showed notable degradation, but mouse spleen samples under the same treatment did not vary substantially from samples stored at 4°C for 6 hr (**Fig. 2**). Replicate sea turtle RBCs samples showed less variation within treatments for fragment size than for DNA yield (**Fig. S5a**,**b**).

Mouse muscle, fish muscle, and fish ovary samples showed considerable accumulation of smaller fragment sizes after one week at room temperature, where blood or muscle samples from other species did not show as dramatic an impact (**Figs. 2, S2, and S3**). However, fish muscle and ovary samples stored at room temperature for just one day still retained high proportions of uHMW DNA with marginal degradation (**Fig. S2**). For mouse muscle, DMSO, EtOH, or DNAgard did not seem to provide any added DNA protection against room temperature conditions (**Figs. 2 and S3**). At the same temperature conditions, mouse samples in Allprotect retained a non-negligible fraction of uHMW DNA, though with some degradation (**Figs. 2 and S3**). Overall, similar to the 4°C exposure, room temperature DMSO and EtOH samples performed relatively well, albeit showing some signs of degradation. Surprisingly, two samples left at room temperature for one week without any preservative (sea turtle RBCs and frog blood) were quite stable and yielded an appreciable fraction of uHMW DNA (**Fig. 2**). Additionally, sea turtle RBCs samples, when preserved with EtOH or even DNAgard and stored at room temperature for 5 months, yielded a large fraction of workable uHMW DNA (**Fig. 2**). This suggested that turtle RBCs may be viable for longer durations at room temperature. Additional replicates and further experimentation will be necessary to determine if the isolated RBCs tissue type or some biological difference in turtles is the key to this stability.

### DNA yield

When the variables were tested individually, vertebrate group explained the least variance in DNA yield (3.69%, DF = 4, F = 3.25, p = 0.01; **Fig. 3d**); temperature treatment explained a similarly small proportion (7.35%, DF = 9, F = 2.88, p = 4.25e-3); preservative explained slightly more of the total variance (10.24%, DF = 6, F = 6.01, p = 1.73e-5; **Fig. 3e**); and tissue type explained the largest amount of variance (46.35%, DF = 5, F = 32.65, p = 2.20e-16; **Fig. 3f**). Both preservative and tissue type together explained 56.59% of the total variance (**Table S2**). Specifically, whole blood tended to generate the highest DNA yields, followed by spleen, RBCs, whole-body, and ovary, while muscle generated relatively lower yield (**Fig. 3f**). In post-hoc tests, whole blood, RBCs, and ovary significantly outperformed muscle (vs. whole blood: t = 11.75, p = 0.002; vs. RBCs: t = 8.36, p < 0.001; vs. ovary: t = 3.28, p = 0.01), while the differences between muscle and whole body or spleen were not significant. Whole blood and RBCs also showed significantly higher yields than ovary samples (vs whole blood: t = 3.89, p = 0.002; vs. RBCs: t = 3.36, p = 0.01). Post-hoc comparisons of different temperature treatments or preservation reagents were not significant, possibly due to the higher variance influenced by the other variables of tissue type and species (**Fig. 3d-f**). Birds tended to have slightly better yields, with a marginally significant effect over non-avian reptiles (t = 3.04, p = 0.02).

### Hi-C sequencing

The VGP is currently using Hi-C reads as a standard tool to generate chromosomal scale assemblies [4,21], as well as to phase haplotypes in some cases [22]. These chromosome interactions are captured *in situ* in the tissue before DNA is isolated and sequencing libraries made. To enable appropriate collection recommendations for use in this technology, we also explore the effect of tissue preservation on the quality of the Hi-C library preparation. Using a single species (zebra finch) we test a subset of tissue preservation methods (flash-frozen, 6 hr at 4°C, one week at room temperature) and tissue types (muscle, blood), with two replicates per treatment combination. These were processed to generate *in situ* Hi-C chromatin interactions maps against the VGP male reference genome [23,24].

We found that blood samples flash-frozen in EtOH yielded similar results compared to our flash-frozen positive control with no added preservative: 75–80% of all read-pairs were derived from *cis* interactions within the same chromosomes (**Fig. 4a**), and among them ∼55–60% were derived from long-range (>15 kb) *cis* interactions. This indicates a high degree of useful long-range intra-chromosomal signal necessary for genome assembly. However, storage of blood in DNAgard resulted in the elimination of almost all *cis* interactions, down to ∼10% total, across temperature treatments (**Fig. 4a-c**), indicating largely random ligations and the loss of useful signal. Blood refrigerated for 6 hr maintained a high yield of long *cis* interactions, both when stored in EtOH and with no preservative. Blood samples stored at one week at room temperature in EtOH also yielded mostly long *cis* interactions similar to the flash-frozen treatments.

**Figure 4.**
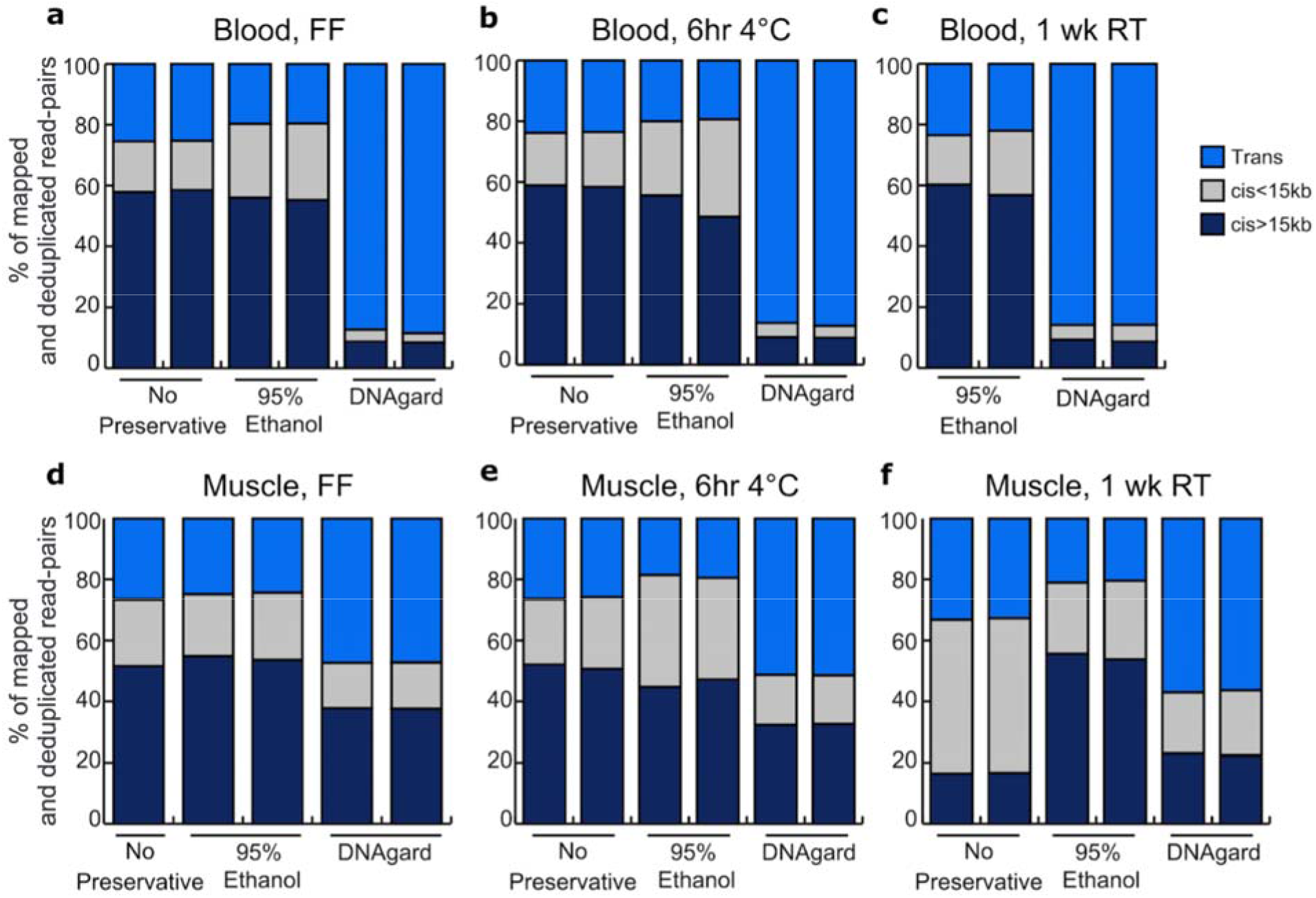
Hi-C platform benchmarking of bird samples. Stacked bar plots denoting proportions of Hi-C reads mapped to the zebra finch reference genome involving different chromosomes (*trans*), on the same chromosome but less than 15 kb apart (*cis* < 15 kb), and on the same chromosome and greater than 15 kb apart (*cis* > 15 kb). Tested samples include blood samples (**a**-**c**), and muscle samples (**d**-**f**). The desirable outcome is to have much greater proportions of Hi-C reads being long-range *cis* pairs, which reflects an efficient capture of long-range interactions needed for genome scaffolding and haplotype phasing. Hi-C data was generated by Arima Genomics following their standard protocol.

Overall, muscle and blood samples performed similarly across all treatments measured using Hi-C reads. They both yielded large amounts of long *cis* interactions (>15 kb) when flash-frozen or refrigerated at 4°C with no preservative or with EtOH (**Fig. 4a-b, d-e**). Muscle and blood samples also responded similarly to preservative treatments, with EtOH samples performing well across treatments and DNAgard samples underperforming across treatments (**Fig. 4**).

## Discussion

During development of the assembly pipeline for the first set of VGP genomes [4], we tested various HMW and uHMW DNA extraction protocols compatible with several LR technologies, including the Qiagen MagAttract HMW DNA, the phenol-chloroform method [5], and the agarose plug protocol. The agarose plug method optimized by Bionano Genomics [7] was the most consistent method for producing a high yield of uHMW DNA suitable across all the LR technologies in the VGP pipeline. This method used agarose as a protective matrix to minimize DNA shearing during the extraction process and had long been shown to be an effective method for isolating megabase-size DNA from organisms including plants, animals, algae, and microbes [7]. In this study, we use only the agarose plug DNA extraction method.

Our study explored the effects of three variables –preservation method, tissue type, and storage temperature– in preserving the high-quality DNA required for generating chromosome-scale genome assemblies in six species representing five major vertebrate lineages. The results identified promising alternatives to the standard flash-freezing method that is not easily performed in the field, particularly the preservation of samples in 95% ethanol (EtOH) or 20– 25% DMSO-EDTA (DMSO) at 4°C.

We did not test all possible combinations of variables, which would require over 252 tests per species, but focused instead on the salient combinations of tissue types, reagents, and protocols that reflect real-world applications. There are also likely intervening stages of exposure to different temperatures, such as immediately post-mortem, that may have a considerable effect in hotter climates and are not simulated here. Despite these limitations, our results are consistent with samples from the over 136 species we have processed for the VGP to date (NCBI Bioproject PRJNA489243 as of July 13, 2021). We believe that the results presented here can inform the many logistical decisions of field researchers collecting samples from wild populations (**Fig. 5**).

**Figure 5.**
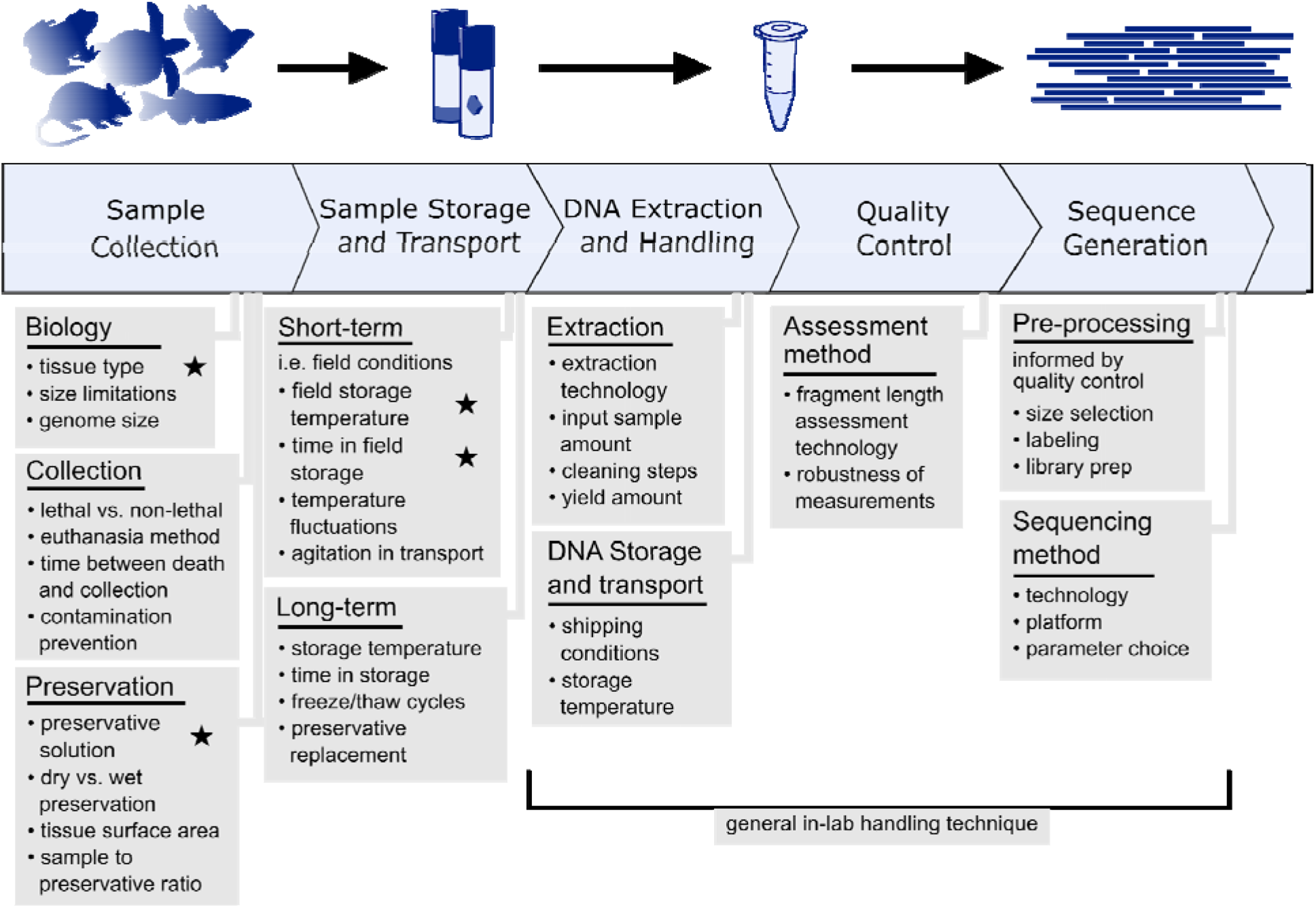
Considerations for collection of tissues for long-read sequencing of non-model organisms. General representation of a sequencing pipeline and considerations that may directly or indirectly affect the quality of sequencing output. Stars indicate particular sources of variation manipulated in this study. Several logistical aspects need to be considered prior to sample collection for uHMW DNA isolation with the goal of producing reference-quality genomes. The collector needs to identify what tissue types can be collected from the target species, what preservation methods and cold storage are available, and how quickly samples can be transported to a –80°C ultra-cold freezer.

Temperature exposure was the strongest predictor of fragment length distribution for these data. The potential of increased temperatures to destabilize DNA is well known, and samples exposed to higher temperatures for a longer period will allow for enzymatic activity that degrades DNA [25]. However, under certain conditions some samples stored at 4°C or even at room temperature show surprising viability. For example, samples preserved in EtOH and refrigerated for up to one week were nearly as good as flash-frozen samples. This is evident through high proportions of uHMW DNA molecules, though with some signs of degradation and variability across species and tissue types.

The ambient temperature of the intended collecting locality should be a major consideration in planning field collections for high-quality samples. Here we test a limited number of samples at 37°C to resemble fieldwork conditions in warmer climates, resulting in no retention of workable amounts of uHMW DNA in any of these samples. Thus, in hotter climates sample cooling or exploring alternative preservatives is critical. Options such as insulated boxes, ice packs, wet ice, dry ice, and electronic coolers should be considered for maintaining samples at low temperatures in the field. To minimize the time before storing in ultra-cold freezers, investigators might also choose to ship samples from the field to the lab before the conclusion of fieldwork. Further experimentation in conditions resembling warmer climates can more precisely define tolerable exposure intervals for sampling targeting uHMW DNA.

The “gold standard” for preserving samples for uHMW DNA extraction remains flash-freezing in liquid nitrogen before ultra-cold storage [9–14]. Our results highlight alternative preservation methods that are more readily available in the field. Liquid nitrogen can be challenging to acquire, contain, and transport in many fieldwork settings. Fortunately, samples preserved in EtOH or DMSO perform well with simple refrigeration. Although a small portion of DMSO samples failed (near-zero DNA extracted) for unclear reasons. In addition, these solutions consistently outperform the commercial preservatives RNAlater and DNAgard. Further, DNAgard is not suitable for maintaining long interaction distances for Hi-C library preparation. While these commercial reagents rely on mechanisms that were likely optimized for preserving lower molecular weight nucleic acids, they appear to be harmful to uHMW DNA and chromosomal 3D interactions. Preservatives that promote cell lysis may undermine the stability of DNA if they cannot adequately counter the increased exposure to sources of chemical degradation [14,25,26]. Of the three commercial reagents tested, Allprotect shows the most promising results for preserving uHMW DNA, but more testing is necessary to better evaluate its performance relative to other preservatives and assess its compatibility with LR technologies.

In addition to popular commercial reagents, we evaluate some of the more commonly applied preservation methods today. EtOH has long been used for preserving samples for DNA analysis, and its proficiency at stabilizing specimens continues to be validated [12,18,27,28]. For example, Mulcahy et al. (2013) studied preservative effects on DNA integrity in white perch and blue crab muscle samples, using only a maximum of 45 kb DNA size resolution. Nevertheless, their finding that EtOH generally performs well as a DNA preservative agent is consistent with our results at this DNA size range. While EtOH is a compelling option, it comes with its own logistical considerations. EtOH can be problematic to transport on commercial flights or trains, or to ship in large quantities. Alternatively, DMSO benefits from fewer transport restrictions, but requires laboratory preparation prior to fieldwork and can be hazardous to handle. Commercial preservation reagents are usually more costly than EtOH or DMSO solutions, but are also under less restricted transport regulations.

The negative impact of DNAgard on Hi-C long-distance *cis* interactions is striking. This solution likely permeates the cell to inhibit nuclease activity, potentially affecting other protein integrity and impeding cross-linking. The increased fraction of inter-chromosomal interactions and decreased fraction of cis-interactions (> 15 kb) together are evidence of DNA degradation. These inter-chromosomal interactions are counter-productive noise with regard to chromosome-level scaffolding in that they erroneously provide scaffolding links between contigs derived from two different chromosomes. Our Hi-C data analysis also indicates, at least for birds, that EtOH storage of blood at 4°C or room temperature for one week or less tends to yield high-quality Hi-C chromatin interaction maps. Excluding samples in DNAgard, blood seems to be slightly more resistant to reducing chromosome interactions than muscle when stored at 4°C or room temperature for one week, which would be a valuable feature for field collection.

Contrary to the differences in Hi-C performance, we did not find notable differences in DNA fragment length distributions between most tissue types. The exception is whole-body fish samples that were all significantly degraded, regardless of treatment. Potentially, this could owe to the larger mass of tissue taking longer to freeze through or infuse with preservative, hence allowing more time for degradation. However, we did observe substantial differences in total DNA yield, where blood and spleen samples tend to yield a larger amount of DNA while muscle samples produce the least. The comparatively lower DNA yield makes muscle samples a less practical choice in species where nucleated blood is available. Lower yield could also be costlier and more time consuming in the long run, as more DNA extractions would be required to achieve the necessary input amount. For species without nucleated blood (mammals), soft tissue samples such as the spleen outperform muscle in terms of yield. Note that low yield does not necessarily preclude muscle samples from usefulness, especially given they still perform well in terms of fragment length if appropriately collected and stored. We note that, as we demonstrated in a related study [29], blood is often not suitable for uHMW mitochondrial DNA extraction, while muscle tends to yield abundant mitochondrial DNA. This is an important consideration if the goal of collection is to sequence the mitochondrial genome.

Our study considers today’s LR sequencing technologies and current DNA isolation protocols. Time will likely continue to yield new methods for preventing, assessing, and mitigating DNA degradation. Even since the outset of this study, promising new extraction methods have become available for uHMW DNA, such as Nanobind DNA extraction (Circulomics, Baltimore, MD, USA). Our comparisons focus on maximizing the quality of field-collected input material and we expect this to be largely independent of downstream extraction methods. Our results and experience acquired with uHMW DNA and Hi-C data for more than 136 VGP genomes produced, yield guidelines for tissue type, preservatives, temperature, and other treatments necessary for generating high-quality genome assemblies from several vertebrate lineages, for laboratory and field collected samples (**Table 1**).

**Table 1:**
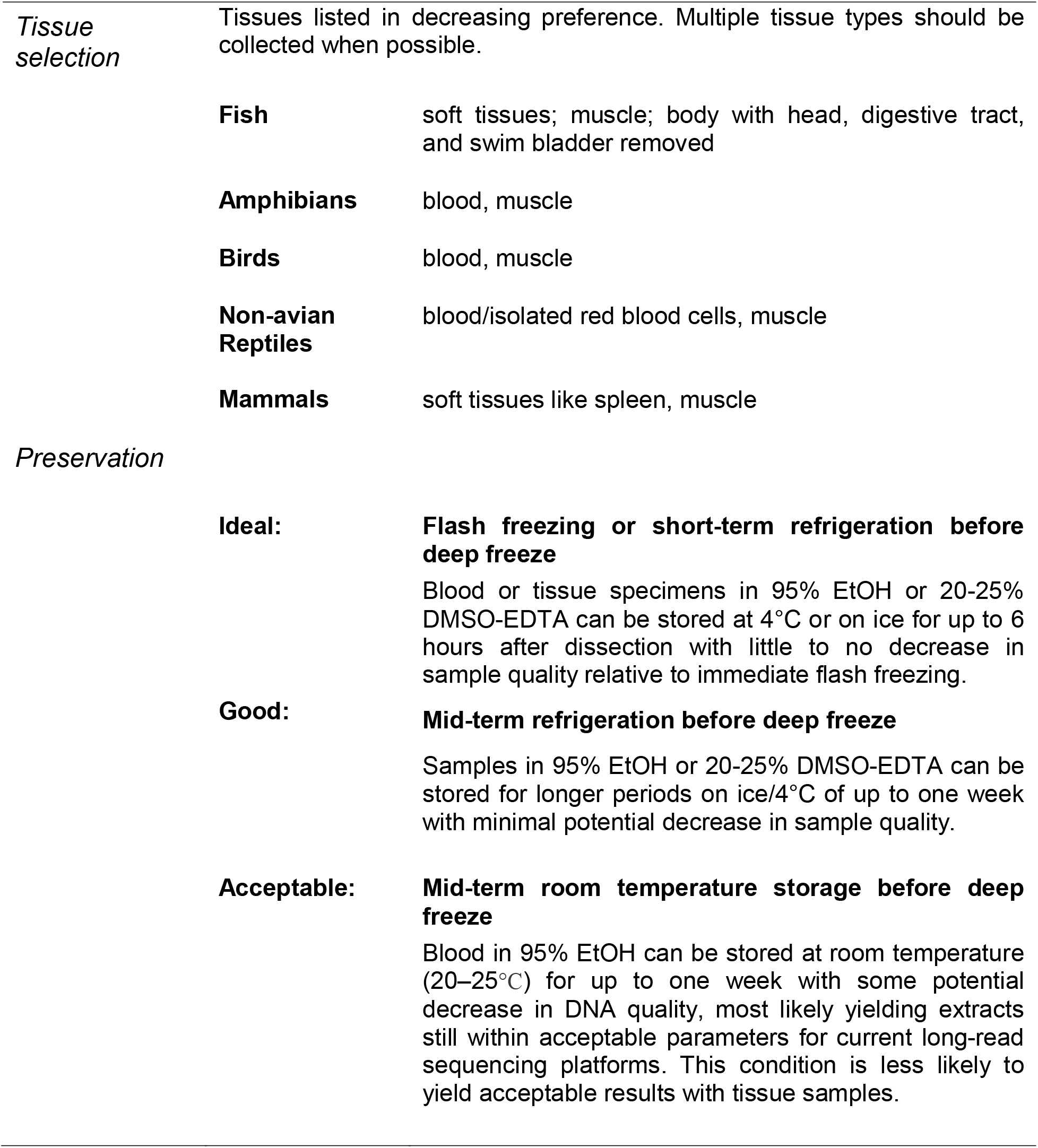
Sample collection guidelines for generating high-quality genomes. Compiled here are guidelines based on the best-performing protocols tested in this study and broadly in the Phase 1 VGP genomes.

|  |  |  |
| --- | --- | --- |
| *Tissue selection* | Tissues listed in decreasing preference. Multiple tissue types should be collected when possible. |  |
|  | **Fish** | soft tissues; muscle; body with head, digestive tract, and swim bladder removed |
|  | **Amphibians** | blood, muscle |
|  | **Birds** | blood, muscle |
|  | **Non-avian Reptiles** | blood/isolated red blood cells, muscle |
|  | **Mammals** | soft tissues like spleen, muscle |
| *Preservation* |  |  |
|  | **Ideal:** | **Flash freezing or short-term refrigeration before deep freeze**<br>Blood or tissue specimens in 95% EtOH or 20-25% DMSO-EDTA can be stored at 4°C or on ice for up to 6 hours after dissection with little to no decrease in sample quality relative to immediate flash freezing. |
|  | **Good:** | **Mid-term refrigeration before deep freeze**<br>Samples in 95% EtOH or 20-25% DMSO-EDTA can be stored for longer periods on ice/4°C of up to one week with minimal potential decrease in sample quality. |
|  | **Acceptable:** | **Mid-term room temperature storage before deep freeze**<br>Blood in 95% EtOH can be stored at room temperature (20–25°C) for up to one week with some potential decrease in DNA quality, most likely yielding extracts still within acceptable parameters for current long-read sequencing platforms. This condition is less likely to yield acceptable results with tissue samples. |

In planning biobanking for genomic purposes, another important strategy is to avoid or reduce the need for field-preserved samples. Seeking out animals already in captive collections and salvaging material reduces the methodological difficulty of preserving samples. Delaying blood collection, biopsy, or euthanasia of wild-caught specimens can also buy researchers time to move into more amenable preservation conditions such as a field station. However, this poses ethical challenges in the care of animals being held for days or weeks, and it is not feasible for larger animals.

Few studies have explored the effects of preservation methods on uHMW DNA integrity [17], but none that we are aware of have done so in as broad a set of field-relevant conditions as in the present study. Being able to collect samples well-suited for producing high-quality genome assemblies is a major undertaking. Our recommendations will enable many new high-quality sample collections and contribute to establishing a greater and more diverse array of vertebrate genomes from around the world.

## Methods

### Sample collection

We collected samples from species representing major taxonomic classes of vertebrates, i.e. house mouse (*Mus musculus*), zebra finch (*Taeniopygia guttata*), Kemp’s Ridley sea turtle (*Lepidochelys kempii*), painted turtle (*Chrysemys picta*), American bullfrog (*Rana catesbeiana*), and zebrafish (*Danio rerio*). All animal handling and euthanasia protocols were approved by the Institutional Animal Care and Use Committees or equivalent regulatory bodies at the respective facilities: The Rockefeller University for the frog and bird samples; the Max Planck Institute for the mouse samples; the University of Toronto for the painted turtle samples; the Wellcome Sanger Institute for the fish samples; and the New England Aquarium rehabilitation facility for the sea turtle samples (**Table S1)**.

For this experiment, tissue samples were collected as available at facilities already handling the target species (**Fig. 1**). The tissue types collected per species are as follows: mouse, spleen and muscle; zebra finch, whole blood and muscle; sea turtle, isolated red blood cells (RBCs); painted turtle, whole blood and muscle; bullfrog, whole blood and muscle; zebrafish, whole body, ovary, and muscle. For all species except the sea turtle and the fish, samples originate from a single individual. In the sea turtle set, duplicate samples were obtained from three individuals. In the fish set tissue samples in some cases originated from different individuals, as their small body size does not allow for sufficient amounts of tissue from a single specimen.

Each taxon required a slightly different handling procedure. All samples except for those from sea turtles were sourced from captive individuals humanely euthanized in a laboratory setting with approved protocols cited below. All soft or fibrous tissue samples were collected in small 20–30 mg pieces until each 2 mL tube had roughly 50–100 mg total to allow for full penetration of the preservative. Mice were euthanized by CO_2_ treatment in a GasDocUnit (Medres Medical Research GmbH, Cologne, Germany) following the instructions of the manufacturer (DD24.1-5131/451/8, Landesdirektion Sachsen). Skeletal muscle and spleen samples were then dissected and placed in standard cryotubes. Birds were euthanized via isoflurane overdose, and whole blood was collected into chilled sodium heparin-treated 1.5 ml microfuge tubes (IACUC #19101-H). Then 25–50 µL was immediately aliquoted into cryotubes. Sea turtle RBCs samples were collected from wild individuals undergoing medical treatment by drawing whole blood into 2 mL sodium heparin-treated collection tubes and then spinning down to separate RBCs from plasma. RBCs were then aliquoted into sodium heparin-treated tubes. Painted turtle samples were collected from one individual euthanized via decapitation as part of another study (AUP 20012070). Painted turtle muscle samples were immediately taken from the pectoral girdle and whole blood was drawn from the heart before placement in standard cryotubes. Frog samples were sourced from one captive adult purchased from Rana Ranch in Twin Falls, Idaho, USA. The frog was euthanized using an intracoelomic injection with Euthasol™ or Fatal-Plus™ (pentobarbital and phenytoin) at a dosage of 100 mg/kg. After confirming that a deep plane of anesthesia was reached, the frog was rapidly and doubly pithed cranially and spinally, then decapitated (19085-USDA). Frog muscle tissue samples were immediately taken from the rear legs and blood was drawn from internal veins before placement in standard cryotubes. We extracted fish samples from multiple lab-raised individuals. To euthanize the fish, we used tricaine and then the brain was destroyed with a scalpel (PPL No.70/7606). We collected white muscle and ovary samples which were dissected out and placed into 2 ml cryotubes immediately after euthanasia. Fish whole-body samples were taken by removing the head, intestines, and swim bladder of individual fish and placing the remaining tissue into a cryotube.

### Preservation treatments

A total of 140 freshly collected samples were subjected to different preservation and temperature treatments to test common preservation methods under simulated field or lab conditions (**Fig. 1**), with flash-frozen samples being used as baseline controls. Preservation method treatments refer to the preservative agent applied directly to the sample before ultra-cold (–80°C) storage; temperature treatments refer to the temperature exposed and the amount of time the sample remained at that temperature before ultra-cold storage.

All temperature treatments were applied immediately upon dissection of the material and placement into specimen tubes. Samples were exposed to temperature treatments of varying lengths of time in refrigeration (4°C), room temperature (20–25°C), and elevated temperature in an incubator to simulate field conditions in a tropical climate (∼37°C). All temperature conditions tested and the samples to which they were applied are as follows: control condition submerged in liquid nitrogen from dissection to ultra-cold storage (all tissue types and species), 6 hr at 4°C (frog blood and muscle, bird blood and muscle, painted turtle blood and muscle, sea turtle RBCs), 16 hr at 4°C (mouse spleen, fish whole body), 1 day at 4°C (fish ovary), 1 week at 4°C (mouse muscle, frog blood and muscle, bird blood and muscle, painted turtle blood and muscle), 1 day at room temperature (fish muscle and ovary), 1 week at room temperature (mouse muscle, frog blood and muscle, bird blood and muscle, painted turtle blood and muscle, sea turtle RBCs, fish muscle and ovary), 4 weeks at room temperature (fish muscle and ovary), 5 months at room temperature (sea turtle RBCs), and 1 week at 37°C (mouse muscle). Storage time at –80°C after treatment and before DNA extraction varied slightly between samples, but such variation is expected to have a negligible impact on sample quality.

The preservation methods tested here include flash-freezing in liquid nitrogen, no added preservative agent, 95% EtOH, 20–25% DMSO-EDTA (DMSO), DNAgard tissue and cells (DNAgard; cat. no. #62001-046, Biomatrica), Allprotect Tissue Reagent (Allprotect; cat. no. 76405, Qiagen), and RNAlater Stabilization Solution (RNAlater; cat. no. AM7021, Invitrogen). Our DMSO recipe was 20–25% DMSO, 25% 0.5 M EDTA, remaining 50–55% H2O, saturated with NaCl. Flash-freezing, EtOH, and DNAgard were tested on all included species and tissue types. DMSO was tested on all species and tissue types except sea turtle RBCs. No-preservative treatments were tested on bullfrog blood, bird blood, painted turtle blood, and sea turtle RBCs. Allprotect was tested on mouse spleen and muscle and fish body. RNAlater was tested on fish ovary and muscle samples.

To gain insights into variation within these treatments, isolated RBCs samples were collected from three different sea turtle individuals and processed separately as biological and technical replicates. The third replicate had insufficient material to test all treatments.

### DNA extraction

We extracted DNA from all tissue samples using the agarose plug protocol as below at VGP data production hubs at the Rockefeller University, Wellcome Sanger Institute, and MPGI Max Planck Institute Dresden (**Table S1**). This method was established, at the time of this experiment, as standard protocol for long-read sequencing in all VGP projects [4]. From each tissue sample, a 30–40 mg piece was weighed and then processed using the Bionano Prep^™^ Animal Tissue DNA Isolation Fibrous Tissue Protocol (Bionano document number 30071) and Soft Tissue Protocol (Bionano document number 30077). Briefly, the fibrous tissue (muscle, whole) pieces were further cut into 3 mm pieces and fixed with 2% formaldehyde and Bionano Prep Animal Tissue Homogenization Buffer. Tissue was blended into a homogenate with a Qiagen Rotor-Stator homogenizer and embedded in 2% agarose plugs cooled to 43°C. Plugs were treated with Proteinase K and RNase A, and washed with 1X Bionano Prep Wash Buffer and 1X TE Buffer (pH 8.0). DNA was recovered with 2 μl of 0.5 U/μl Agarase enzyme per plug for 45 minutes at 43°C and further purified by drop dialysis with 1X TE Buffer. The soft tissue (spleen, ovary) pieces were further cut into 3 mm pieces and then homogenized with a tissue grinder followed by a DNA stabilization step with ethanol. The homogenate pellet was then embedded in 2% agarose plugs as in the fibrous tissue protocol above. For blood samples, DNA was extracted from whole blood or RBCs following the unpublished Bionano Frozen Whole Nucleated Blood Stored in Ethanol – DNA Isolation Guidelines. The ethanol supernatant was removed and the blood pellet was resuspended in Bionano Cell Buffer in a 1:2 dilution. For samples that freeze solidly at –80°C, tubes were thawed at 37°C for 2–4 minutes. The same Bionano guidelines for nucleated blood in ethanol were modified by adding 1–2 additional centrifugation steps at 5,000X g for 10 min prior to removing DNAgard supernatant and homogenizing blood cells in Bionano Cell Buffer in a 1:2 dilution. All samples were mixed with 36 µl agarose and placed in plug molds following the animal tissue protocol.

### Assessing sample purity and yield

All extractions had sufficient DNA yield to measure except one: mouse spleen tissue in DMSO. This sample congealed and solidified in such a way that no DNA could be extracted. To measure DNA yield and purity, we used both the fluorescence-based Broad Range Qubit® assay and absorbance-based Nanodrop One^™^. To measure yield, 2 μl aliquots of gDNA were taken from the top, middle, and bottom of each DNA sample and diluted in a Qubit Working Solution of 1:200 Dye Assay Reagent with BR Dilution Buffer. Sample concentrations were recorded on a Qubit 4 Fluorometer. The concentration of the top, middle, and bottom readings were averaged to estimate the concentration of each DNA sample. Spectrophotometry was then performed on a Nanodrop One to measure sample purity in terms of the 260/230 and 260/280 nm ratios.

### Assessing sample fragment size distributions

Fragment length distributions of samples were measured with at least one of two available methods: Pulsed-field Gel Electrophoresis (PFGE) or the Agilent Femto Pulse system (FEMTO). PFGE was performed using the Sage Science™ Pippin Pulse gel system with the Lambda PFG Ladder (New England Biolabs). To quantify fragment length distribution from PFGE gel images, we compared the proportions of signal above and below 145 kb. This was done using the program ImageJ [30] following Mulcahy et al. (2016) based on the Gel Analysis tool in ImageJ. Further quantifying of the PFGE signal below 145 kb, such as the relative amount of low molecular weight DNA, was not robust due to compression or streaking obscuring smaller fragment patterns. Concise visualization of gel plot profiles was produced in the R package ggridges [31] with a custom Python script for piecewise linear scaling across different gels according to a common size standard. Grey-value intensity measured in ImageJ was scaled locally in each lane and cropped to the gel boundary such that, excluding the well, the brightest value along the lane became 100 and the darkest became 0. Analysis of FEMTO outputs was carried out in the ProSize Data Analysis Software. First, each trace was assessed for signs of an unreliable run, including ladder quality, loading concentration, raised baseline, and unusual smear patterns. Runs with these hallmarks were not incorporated further. Because signals above 165 kb are not reliable on FEMTO, we only considered signals within the range of 1.3–165 kb. We then recorded the proportion of the sample measuring above 45 kb. Further visualization of FEMTO traces were made in the same manner as above with a custom python script and the R package ggridges, except scaling to a size standard was done in ProSize. Yields were insufficient for fragment size analysis from frog muscle in DMSO for one week at 4°C and 6 hr at 4°C and mouse spleen in DMSO for 16 hr at 4°C.

### Statistical analysis

We used linear modeling in the R statistical package to explore the relative contribution of several factors to the variance in DNA yield and fragment length among tests. The three response variables (DNA yield per unit mass (yield), PFGE proportion > 145 kb (PFGE), FEMTO proportion > 45 kb (FEMTO)) were modeled separately. The data for each model were samples with those measurements, and all conditions had at least two replicates (yield: n = 139, PFGE: n = 102, FEMTO: n = 108). DNA yield was log-transformed using the natural logarithm to satisfy assumptions of normality and modeled with temperature, preservative, vertebrate group, and tissue type included as fixed effects. Homoscedasticity was checked after modeling and found to conform to assumptions. PFGE proportion and FEMTO proportion were modeled with quasibinomial error distributions with temperature, preservation method, and tissue type included as fixed effects. Vertebrate group was not included in the final fragment length models due to collinearity with tissue type. Post-hoc tests were done using the glht function of the R package multicomp to examine differences between the levels of each factor. Further model details including p-values and contingency tables are available in the supplementary materials (**Tables S2 and S3**).

### Hi-C library preparation and sequencing

Because Hi-C methods require intact cell nuclei, we tested a subset of bird samples from our preservation experiments directly using the Arima-HiC platform. We tested blood and muscle samples in three different treatments: without preservatives, in EtOH, and in DNAgard. Each preservation method was subjected to three temperature treatments: immediately flash-frozen, 6 hr at 4°C, and one week at room temperature (20–25°C). After temperature treatment, each sample was moved to –80°C. Blood with no preservative at room temperature for one week was excluded from this set. Two technical replicates of each sample were prepared and sequenced at Arima Genomics following their standard protocol. We measured the performance of Arima-HiC runs by mapping the sequence reads to the zebra finch reference genome (GCA_003957565.1) to determine the proximity of ligated sequence pairs. Assessments were made based on the ratios of *cis* (intra-chromosome) to *trans* (inter-chromosome) read pairs as well as the total percentage comprised of long-distance (> 15 kb) cis pairs.

## Data availability

Sample information, PFGE measurements, FEMTO measurements, and DNA yield data can be found in the supplemental materials. Raw FEMTO outputs, PFGE gel images, and Hi-C read-pairs are available on Dryad (doi:10.5061/dryad.000000041).

## Acknowledgments

This research was supported by Howard Hughes Medical Institute Funds and Rockefeller University Startup funds to EDJ, institutional funds of the Max Planck Institute of Molecular Cell Biology and Genetics, and funds by the Wellcome Trust made out to DNAP R&D team at Wellcome Sanger Institute. IB’s time was supported by Wellcome grants WT207492 and 104640/Z/14/Z, 092096/Z/10/Z. Sampling was facilitated by Leslie Buck and Mouska Patang for the painted turtle and Brian Fabella for the bullfrog. Sea turtle sampling was conducted and generously facilitated by the New England Aquarium and Massachusetts Audubon Wellfleet Bay Wildlife Sanctuary authorized under USFWS permit TE01150C-1; sample transfer to LMK was permitted via a USFWS special authorization letter.

## Author contributions

J.M., S.W., A.F.S., I.B., L.M.K., T.L., A.J.C., R.W.M., A.D.S., P.A.M., E.D.J., and O.F. initially conceptualized the study. H.A.D., J.M., J.B., S.W., A.F.S., S.M., O.V.P., I.B., K.O., M.S., W.T., A.K., L.M.K., E.D.J., and O.F., carried out data collection and preprocessing. H.A.D., J.B., G.F., and A.F.S. analyzed the data and produced the figures. The manuscript was drafted by H.A.D., J.M., J.B., G.F., E.D.J., and O.F., and all authors contributed to revisions.

