## Supplemental Figures for "Benchmarking ultra-high molecular weight DNA preservation methods for long-read and long-range sequencing"

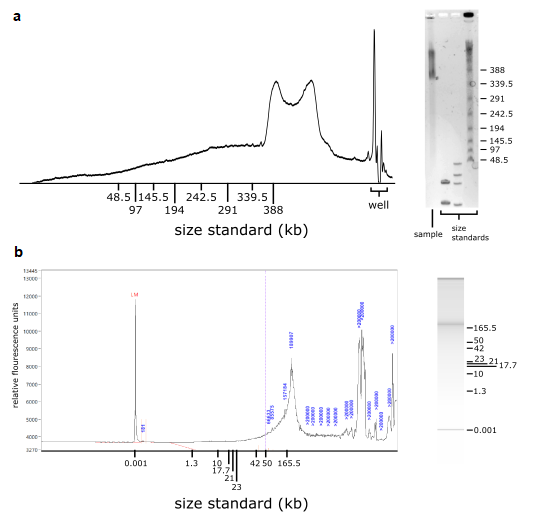

**Figure S1. Example of genomic DNA traces of the same sample made with two different methods.** DNA extract from flash frozen mouse muscle was measured with (**a**) Pulsed-field gel electrophoresis via the software ImageJ and (**b**) Agilent FEMTO Pulse via the software ProSize. Traces are displayed as gel images (right) and as plot profiles (left).

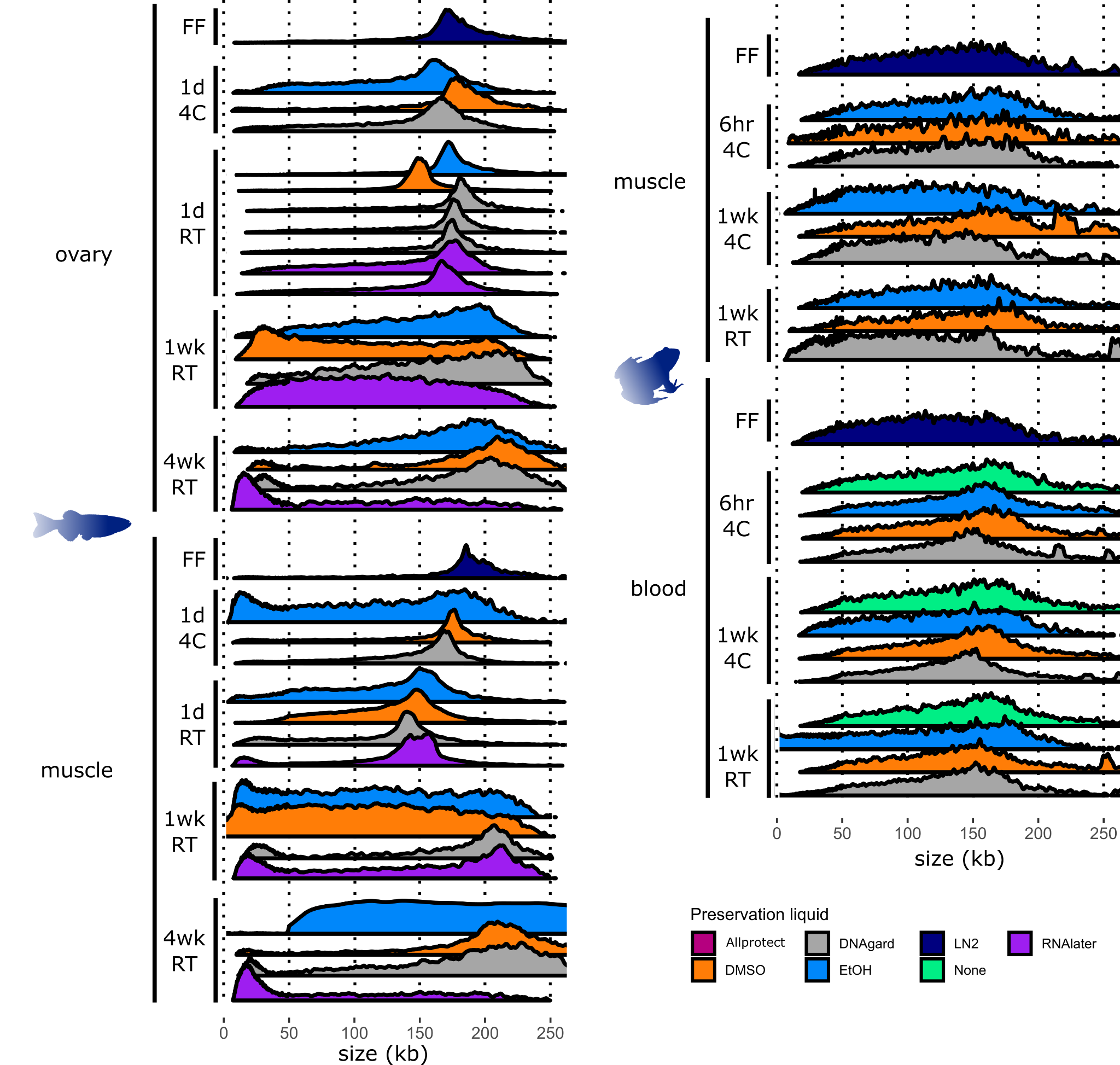

**Figure S2. Plot profiles of FEMTO results on fish and frog samples.** Agilent FEMTO Pulse traces for fish and frog samples are visualized as overlapping ridgeline plots. Each ridgeline plot corresponds to a single sample. X-axis values are scaled via ProSize Data Analysis Software. Y-axis of each plot is scaled independently to ignore peaks outside the range of 10-165kb such that the highest value in that range of each plot becomes 100 and the lowest value becomes 0. Plots with excess noise are generally ones that have low signal.

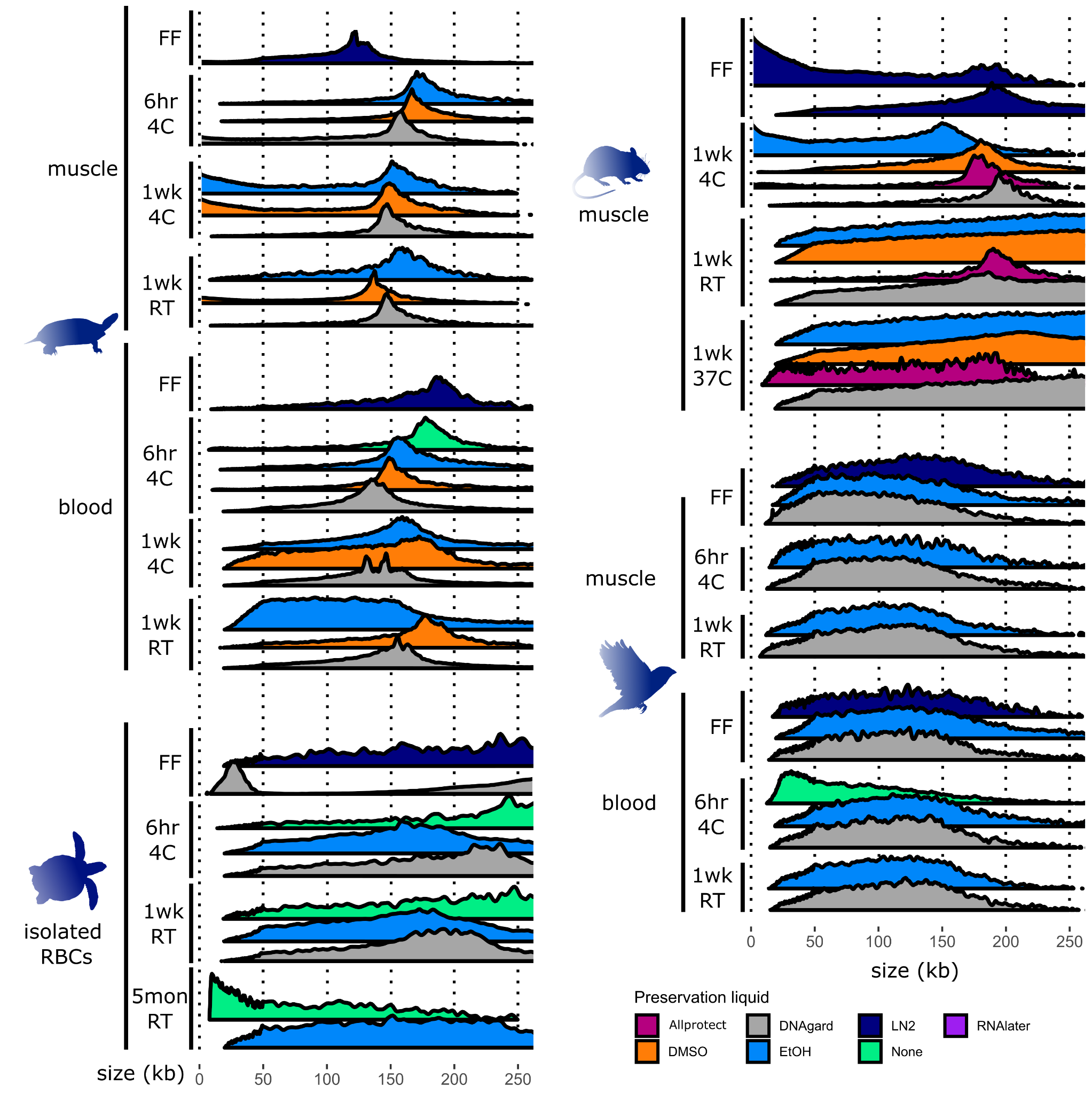

**Figure S3. Plot profiles of FEMTO results on turtle, mouse, and bird samples**. Agilent FEMTO Pulse traces for turtle, mouse, and bird samples are visualized as overlapping ridgeline plots. Each ridgeline plot corresponds to a single sample. X-axis values are scaled via ProSize Data Analysis Software. Y-axis of each plot is scaled independently to ignore peaks outside the range of 10-165kb such that the highest value in that range of each plot becomes 100 and the lowest value becomes 0. Plots with excess noise are generally ones that have low signal.

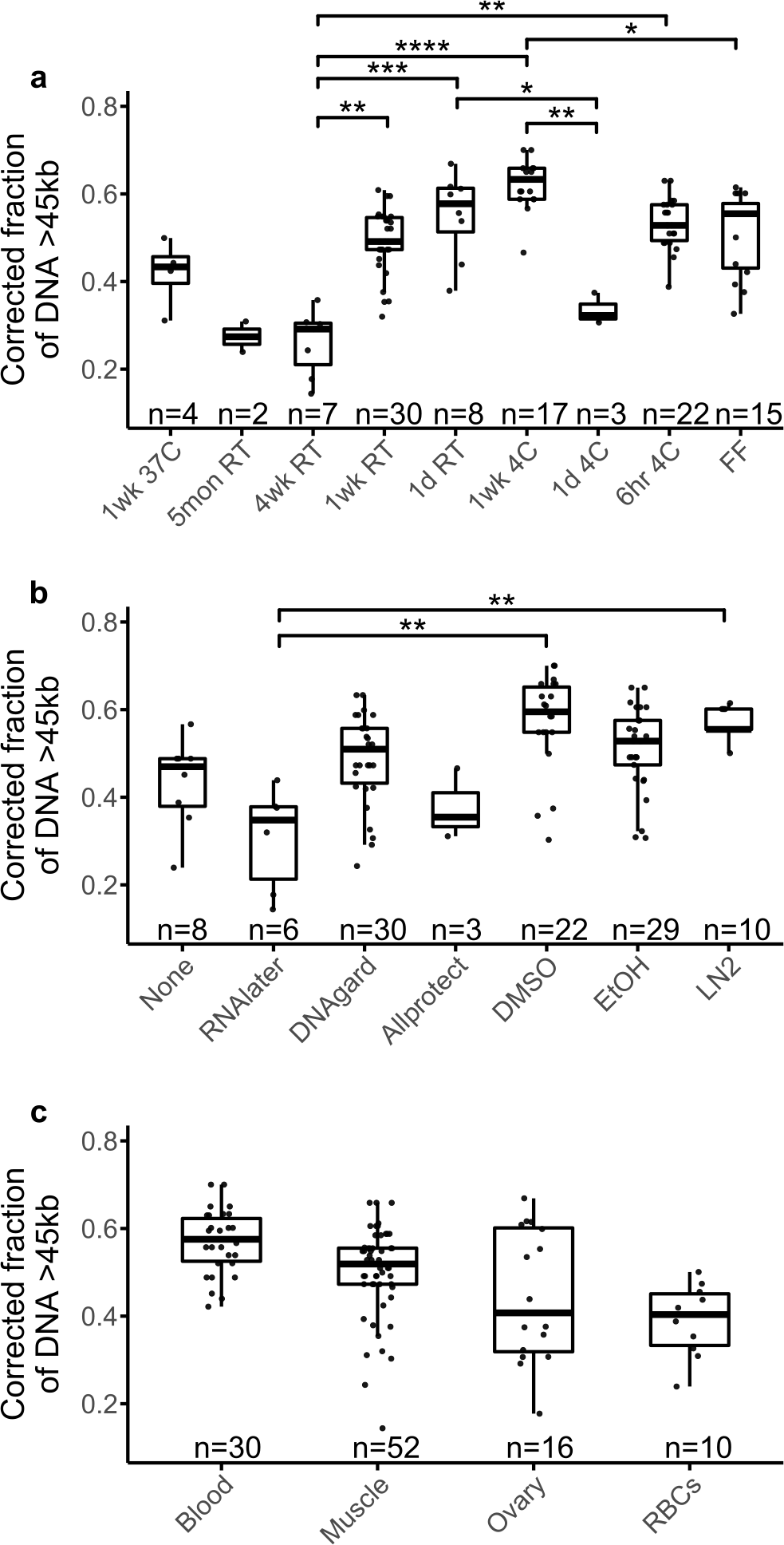

**Figure S4. Testing of different variables on uHMW DNA proportion as measured by FEMTO.** Distributions of sample groups are overlaid with results of linear modeling of fragment length (n = 108). Shown are box-and-whisker plots, with the median, quartiles, and full range of individual observations. Fragment length was quantified here as the proportion of signal between 45kb and 165kb as measured on the Agilent FEMTO Pulse system and modeled in a generalized linear model with temperature (**a**), preservative (**b**), sample type (**c**) as predictors. Significant relationships from post-hoc comparisons are shown as connecting bars with significance levels: **** p < 0.0001, *** p < 0.001, ** p < 0.01, * p < 0.05.

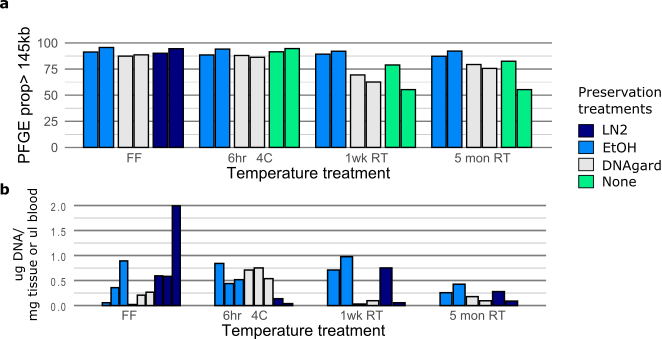

**Figure S5. Comparisons of replicate sea turtle RBCs samples.** (**a**) DNA yields per unit of sample input shown for up to three replicates per treatment. (**b**) Bar plots of PFGE measurements of signal proportion greater than 145 kb, excluding the well. The ordering of sea turtle replicates is plotted consistently from left to right, i.e.- the first bar of each treatment is from the same individual.
